# Brain encoding of naturalistic, continuous, and unpredictable tactile events

**DOI:** 10.1101/2023.12.08.570756

**Authors:** Nicolò Castellani, Alessandra Federici, Marta Fantoni, Emiliano Ricciardi, Francesca Garbarini, Davide Bottari

**Affiliations:** MoMiLab, IMT School for Advanced Studies, Lucca, Italy; Manibus Lab, University of Turin, Turin, Italy

## Abstract

Studies employing EEG to measure somatosensory responses have been typically optimized to compute event-related potentials in response to discrete events (ERPs). However, tactile interactions involve continuous processing of non-stationary inputs that change in location, duration, and intensity. To fill this gap, this study aims to demonstrate the possibility of measuring the neural tracking of continuous and unpredictable tactile information. Twenty-seven young adults (females = 15) were continuously and passively stimulated with a random series of gentle brushes on single fingers of each hand, which were covered from view. Thus, tactile stimulations were unique for each participant, and stimulated fingers. An encoding model measured the degree of synchronization between brain activity and continuous tactile input, generating a temporal response function (TRF). Brain topographies associated with the encoding of each finger stimulation showed a contralateral response at central sensors starting at 50 ms and peaking at about 140 ms of lag, followed by a bilateral response at about 240 ms. A series of analyses highlighted that reliable tactile TRF emerged after just 3 minutes of stimulation. Strikingly, topographical patterns of the TRF allowed discriminating digit lateralization across hands and digit representation within each hand. Our results demonstrated for the first time the possibility of using EEG to measure the neural tracking of a naturalistic, continuous, and unpredictable stimulation in the somatosensory domain. Crucially, this approach allows the study of brain activity following individualized, idiosyncratic tactile events to the fingers.

**Significant Statement:** This study expands the current research conducted on neural tracking, opening the exploration of idiosyncratic tactile events and overcoming constraints of laboratory tasks that typically rely on discrete events. We validated a protocol for the ecological investigations of continuous, slow, tactile processing of the hands. The employed approach enriches the possible use of the EEG to characterize somatosensory neural representations of tactile events. Findings unravel coherent neural responses to continuous and naturalistic touch, with sensitivity for digit lateralization and representation.

## 1. Introduction

EEG is one of the most popular methods to investigate neurophysiological correlates of tactile processing. The majority of studies employing this technique have been optimized for the computation of event-related potentials (ERPs) which require the recurrence of specific events a multitude of times (Fossataro et al., 2023; Galigani et al., 2021; Hogendoorn et al., 2015; Macerollo et al., 2018; Ronga et al., 2021; Ronga et al., 2021b). Following the average of electrical activity across repetitions, the emerging ERP estimates the brain response to the onset of discrete tactile events. ERPs have an excellent signal-to-noise ratio (SNR) and can be easily compared between different experimental conditions. However, in everyday life, touch is experienced differently (Serino & Haggard, 2010). The body-environment interaction entails continuous processing of non-stationary inputs (time-variant) that change in space (skin portion) and quality (type of touch).

Novel methodological approaches have recently emerged, allowing the investigation of electrophysiological data acquired during continuous sensory stimulation (Crosse et al., 2016; Morales & Bowers, 2022). Neuronal populations can synchronize their activity (through alignment of the phase) to temporal profiles of a continuous input (Lakatos et al., 2019; Obleser & Kayser, 2019). This neural tracking has been measured with a variety of stimuli and tasks, including visual stimulation (Bourguignon et al., 2020; Hauswald et al., 2018; King et al., 2016; Spaak et al., 2014), speech processing (Di Liberto et al., 2015; Keitel et al., 2017; Kösem et al., 2018; Zoefel & VanRullen, 2016), music (Zuk et al., 2021) and multisensory inputs (Jessen et al., 2019; Thézé et al., 2020). In the case of the forward or encoding models, the mathematical framework underpinning the neural tracking aims to predict ongoing brain activity from continuous stimulus features (e.g., Crosse et al., 2016; Lakatos et al., 2019). These analysis approaches are increasingly popular in neuroscientific research (Crosse et al., 2016, 2021); however, they are rarely employed in the somatosensory system (Chota et al., 2023; Maallo et al., 2022), and, importantly, naturalistic tactile inputs have never been employed. The possibility of investigating the association between varying somatosensory inputs and ongoing brain activity could represent a leap toward unraveling how the brain represents sensory input.

Here, we demonstrated the possibility of measuring the neural tracking of continuous and unpredictable tactile stimulation of human hand fingers. Each participant was exposed to non-standardized continuous somatosensory stimulations to fingers. To extract the time-varying information of a gentle touch, we recorded sounds generated by tactile stimulations using a miniaturized microphone fixed on a brush. We then extracted the sound envelope, which represented the information concerning the friction between the participants’ skin and the brush. In this case, the sound envelope provided information regarding the onset and offset of the tactile stimulation, its dynamic and frequency.

Since encoding for different sensory stimuli highlighted responses that are neurophysiologically interpretable like ERPs (Crosse et al., 2016), the temporal response function (TRF) derived from the forward model should resemble the somatosensory ERPs in both timing and topographical distribution.

This work aimed to extend and validate the neural tracking approach in the somatosensory domain, possibly opening a new branch of study to measure how the brain encodes naturalistic tactile events. To this aim, we (i) computed the TRF at the individual level based on the specific and unique tactile stimulations each subject received; (ii) estimated a group-level TRF to assess the solidity of the measurement across subjects, despite each of them being stimulated differently; (iii) ensured that we measured a pure tactile TRF, and that spurious visual and auditory input could not explain the observed effects; (iv) evaluated whether the TRF topographies allowed dissociating digit lateralization across the hands and digit representation within each hand; (v) tested the minimum amount of data to reconstruct a solid and robust average TRF associated with continuous and naturalistic tactile stimulation of the hands by comparing it with noise.

Results clearly revealed a successful measurement of how the brain encodes naturalistic, continuous, and unpredictable tactile events.

## 2. Materials and Methods

### 2.1 Participants

Since, to the best of our knowledge, this is the first study assessing neural tracking of naturalistic tactile stimulation, we estimated the sample size allowing to estimate a reliable neural tracking as compared to noise. We focused our analysis on a central contralateral cluster of electrodes (electrodes [E22, E25, E26, E27, E28] for right stimulation and electrodes [E42, E45, E46, E48, E49] for left stimulation), in line with the large literature regarding somatosensory processing (Fossataro et al., 2023; Hogendoorn et al., 2015; Macerollo et al., 2018; Pyasik et al., 2021; Ronga et al., 2021); the frequency of interest [0.5-6 Hz] was defined after the inspection of a spectral power plot (see Figure 1); the time-window for the data simulation was [0.5-200 ms] which comprises the main somatosensory responses starting from the earliest component. We ran a power analysis simulating the main contrast of interest (neural activity measured in experimental conditions vs. control condition; see 2.2) by using a cluster-based permutation test and the data of 10 participants (Wang & Zhang, 2021). The sample size estimation resulted in 27 participants. Thus, we recruited and tested in this study a total of 27 young-adult volunteers (female = 15; mean ± SE: 27.92 ± 0.36 years, range 25-32). They all had normal or corrected to normal vision and were right-handed by self-report; none reported a history of neurological disorders. Participants were naïve to the purpose of the experiment. All participants were informed about the experimental procedure and signed a written informed consent before testing. The study was approved by the regional ethical committee. The study protocol adhered to the guidelines of the Declaration of Helsinki (World Medical Association, 2013).

**Fig. 1.**
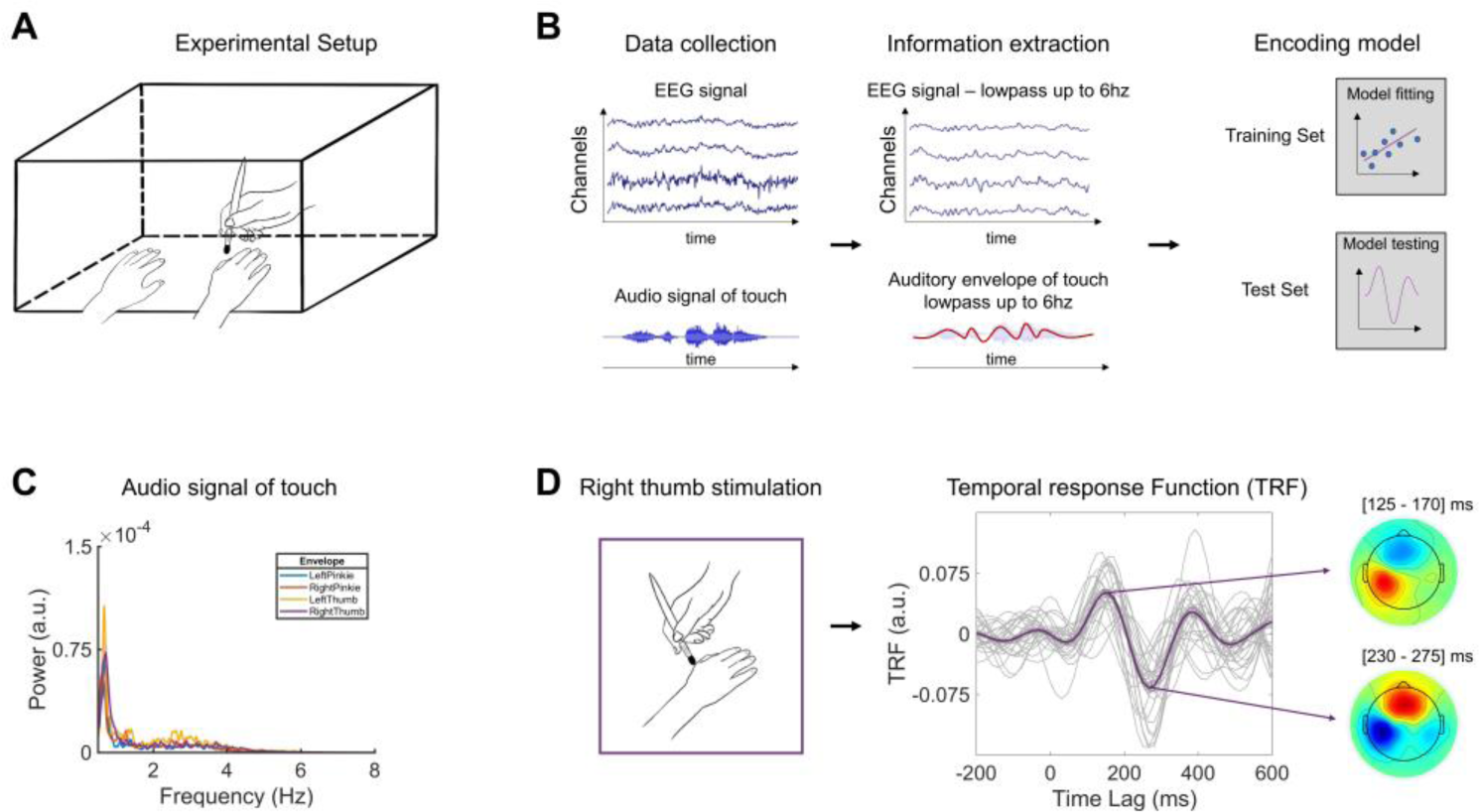
(A) Experimental set-up. The participant’s hands are kept comfortably on the table, covered from view by a wooden box. With an ad-hoc built brush, the experimenter stimulated the participant’s fingers (e.g., the right thumb). (B) Data processing and performed model. The figure summarizes the analysis pipeline. After data collection, slow activity in both EEG signal and auditory information associated with continuous touch was extracted. An encoding model was used to obtain the TRFs. (C) Power spectrum (0.5 – 8 Hz) of the audio envelope of touch for each experimental condition. (D) Tactile TRF results of right thumb stimulation. The grand average TRF (N=27) is depicted in purple, with single participants in light grey. The topographies show the spatial distribution of the grand averaged TRFs of the first positive (∼140ms) and negative (∼245ms) responses in two 45 ms time windows centred on each peak.

### 2.2 Procedure

The participants underwent a (i) preliminary behavioural and an (ii) EEG experiment in a single session. The behavioural experiment always preceded the EEG. For both experiments, participants were seated in a comfortable chair and kept both hands on a table. The experimenter was seated in front of the participant. Crucially, a box covered both participants’ and experimenter’s hands from view (see Figure 1). Experiments were conducted in a soundproof chamber (BOXY, B-Beng s.r.l., Italy).

The preliminary behavioural experiment aimed to evaluate if the subject could perceive auditory information related to the friction between the brush and the skin. The experimenter performed 40 trials: 20 in which he stimulated his own left index finger with a single brush and 20 without stimulation. The order of the trial was fully randomized for each participant. The instructions for the subject were to guess if there was or not a stimulation. The two kinds of trials (i.e., stimulation vs. no stimulation) and the two possible responses (i.e., yes vs. no) derived four possible results: hits (i.e., stimulation and yes), omissions (i.e., stimulation and no), false alarm (i.e., no stimulation and yes) and correct rejection (i.e., no stimulation and no). The whole preliminary experiment lasted for 4 minutes.

During the EEG experiment, participants kept the same posture as in the preliminary one, with their hands hidden from view, and placed comfortably on the table. Participants were instructed to look at a fixation cross positioned at the centre-top of the box, equidistant from their hands, and to pay attention to the tactile stimulation. The main experiment had *five conditions*: right thumb, right pinkie, left thumb, left pinkie and a control condition. In the control condition, everything was kept constant except the target of the tactile stimulation. The experimenter stimulated his own left-hand thumb to rule out the possibility that auditory input (or any other visual cue) associated with brushing participants’ fingers during real tactile stimulation could contribute to neural entrainment. Only the forearms of the experimenter were hidden; thus, it is possible that during brushing, participants could entrain with other movements (shoulder, head, etc.) of the experimenter. Three trials for each condition were performed, resulting in 15 experimental trials lasting three minutes each. During each trial, the experimenter stimulated the specific finger through an ad-hoc built brush, recording the friction between the brush and the skin. Each trial consists of a three-minute continuous and naturalistic stimulation of a specific finger (either the left pinkie, left thumb, right pinkie, or right thumb). The experimenter kept stimulating a specific finger for an entire trial. The brushing direction was constant, from the upper finger back through the nail of the finger. The variability is given by using different spots on the finger as starting points, different velocities, pressures of each specific stroke, and random stroke frequencies inside the trial. Crucially, each trial and condition stimulation was unique (random) for each participant. The total experimental duration was 45 minutes. Participants were encouraged to take small pauses between each trial, during which they moved their hands. The order of experimental conditions was controlled and randomized by PsychoPy3® (v2020.1.3). An external sound was used to code the beginning of each trial and ensure precise tactile stimulation/EEG synchronization.

### 2.3 Behavioural Data analysis

The behavioural data collected in the preliminary experiment: stimulation (present vs. absent) and response (yes vs. no). This resulted in a percentage ratio of hit (yes/present), missing (no/present), false alarm (yes/absent) and correct rejection (no/absent). A within subject paired t-test was performed to contrast hit and missing. Finally, the d’, a measure to assess the sensitivity of the participants’ responses, was computed by subtracting the z-score of the false alarm rate from the z-score of the hit rate.

### 2.4 EEG recordings and preprocessing

The EEG recordings were acquired at a sampling rate of 500 Hz using NetStation5 software and a Net Amps 400 EGI amplifier connected to 64 electrodes HydroCel Geodesic Sensor Net (Electrical Geodesics, Inc.), all signals were referenced to vertex (additional channel E65/Cz). Electrode impedances were kept below 30 kΩ.

We preprocessed continuous EEG raw data offline using MATLAB (R2020b, Mathworks Inc., Natick, MA) and EEGLAB toolbox (version 264 14.1.2b, Delorme & Makeig, 2004). The preprocessing of the raw data followed these steps: (i) band-pass filtered from 1 to 40 Hz (low-pass: FIR filter, filter order: 100, window type: Hann; high-pass: FIR filter, filter order: 500, window type: Hann); (ii) downsampled to 250 Hz; (iii) Independent Component Analysis (ICA) (Jung et al., 2000) to remove artefactual components related to blinks, eye movements and heartbeat. To semi-automatize this process in a data-driven manner, the ICA weights were applied to the raw data, the components were first labelled using ICLabel (Pion-Tonachini et al., 2019). Hence, the most prototypic component for each artefact was selected. It resulted in a total of three prototypic components for eye blink, eye movements and heartbeat, respectively, chosen by the highest probability value given by ICLabel. These three prototypic components were used as templates for the automatic identification of similar components with CORRMAP (Version 1.03; Campos Viola et al., 2009). CORRMAP automatically detected all components with a correlation coefficient exceeding a threshold of 0.8 with the prototypic ones. These identified components were then removed from the original raw data (removed components per participant mean±SE: 3.04±0.22). (iv) Noisy channel detection and spherical interpolation (Interpolated channel mean±SE: 2.37±0.22); (v) re-referenced to standard average reference; (vi) band-pass filtered from 0.5 to 6 Hz (low-pass: FIR filter, filter order: 200, window type: Hann; high-pass: FIR filter, filter order: 2000, window type: Hann); (vii) downsampled to 64Hz; (viii) data epoching according to the onset of the tactile stimulation, considering a delay between the auditory event that coded the beginning of the stimulation and its corresponding EEG marker (i.e., 89 ms), and (ix) segmented into 1 minute long trials, resulting in 9 trials per condition (N=5) per subject (N=27). The stimulation and EEG recording occurred for three minutes in each trial, so the 1-minute segmentation did not aggregate non-consecutive events (Crosse et al., 2021). Lastly, (x) EEG data corresponding to a 1-minute segment were normalized, computing the z-scores to optimize the cross-validation procedure.

### 2.5 Tactile Envelope Extraction

The sound corresponding to the tactile stimulations was registered online through a custom-built brush with a miniaturized condensation microphone (RØDE, Lavalier GO). For each condition, the sounds of the tactile stimulations were first concatenated, leading to 9 minutes of recording, and segmented into one-minute trials, obtaining 9 trials for each condition and subject. We extracted the tactile envelope for each trial by computing (i) the absolute value of the Hilbert transform of the stimulations, (ii) applying a low-pass filter (3rd-order Butterworth filter with 6 Hz as cut-off frequency), (iii) downsampling the signal to 64 Hz (i.e., matching the downsampling of the EEG data, e.g., Mirkovic et al., 2015) and (iv) normalizing them by dividing for the maximum value.

### 2.6 Temporal response function estimation

The temporal response function estimation was performed with the mTRF toolbox (Crosse et al., 2016). This TRF approach allows to build an encoding model to predict an unknown EEG response from the stimuli features (i.e., auditory envelope of touch). This approach has already been employed to evaluate the neural tracking of speech sound, linguistic and visual properties (King et al., 2016; Plass et al., 2020; Thézé et al., 2020). This mathematical function exploits ridge regression. The TRF is a mathematical filter that describes the linear relation of the brain response to the stimuli features in a pre-specified number of time points associated with the stimuli occurrence (Crosse et al., 2016). The ridge regression is used to avoid the problem of overfitting the model (i.e., building a model that perfectly predicts the data used in the training set but fails to predict unseen data). This function relies on the ridge parameter, which prevents overfitting by penalizing the model weights to keep the model as generalizable as possible. To avoid model overfitting, the optimal regularization parameter was empirically identified, employing a leave-one-out cross-validation procedure, choosing a predetermined selection of ridge values (i.e., from 10^-6^ to 10^8^) for time lags from -400 to 600 ms. To choose the regularization parameter, the mean squared error (MSE) between the real and the predicted EEG response, for each participant and condition, was evaluated. Critically, the parameter yielding the lowest MSE on the majority of test datasets was selected. The optimal parameter (λ=10^2^) was selected and kept constant across participants and experimental conditions to generalize the procedure at the group level. It is important to underline, that despite gaining generalizability, this may lead to suboptimal TRF estimation for some individual subjects. We computed independent encoding models for each EEG channel (N=64) in each participant for each condition. We employed the data in a window starting from -400 to 600 ms. The tactile envelope was extracted below 6 Hz since these frequencies were the most represented in the tactile stimulation (see Figure 1). The encoding models were trained employing the tactile envelope as predictor, using a leave-one-out cross-validation procedure (Crosse et al., 2016). Crucially, the TRF model for each participant is the average of each model computed for every left-out-trial. Finally, the grand average TRF models were computed by averaging TRFs across participants within conditions. The main advantage of the TRF model is that encoding model weights are physiologically interpretable (Crosse et al., 2016; Haufe et al., 2014). These weights, computed for each EEG channel, allow to explore the topography of the TRFs (similar to the ERP response) both regarding amplitude and directionality of the data.

### 2.7 Statistical analysis

First, for each condition we evaluated the existence of somatosensory entrainment against the noise of the data. The spatiotemporal profiles of each experimental condition TRF (*tactile TRF*) was contrasted to a null effect (*null TRF*) obtained by 100 permutations of the original data. Statistical comparisons were conducted employing paired t-test with a cluster-based permutation approach (Maris & Oostenveld, 2007, Monte Carlo method 1000 permutations, the cluster-based p-value was 0.025, accounting for two-tailed testing; from now on significant results are labeled as p_clust_ < 0.05) across 0 to 400 ms time-lags and all electrodes. Cluster-based permutation statistics allowed to control for the multiple comparisons problem associated with testing across time lags and sensors. The summed t-statistics associated with the comparison of interest are clustered in time and space and contrasted versus the summed t-statistics associated with the comparison following a random partition obtained by shuffling the experimental condition labels (i.e., experimental TRF vs. *null TRF*).

Subsequently, to test the robustness of the somatosensory entrainment, we contrasted the spatiotemporal profiles of *tactile TRF* (i.e., real touch) against the control condition (*control TRF*). Namely, we contrasted each experimental condition (i.e., left thumb, left pinkie, right thumb, and right pinkie) versus the control condition in which the experimenter’s hand was stimulated (see Method, 2.2 procedure). These contrasts allowed to firmly exclude the possibility that spurious non-somatosensory cues (e.g., auditory and visual information) could explain the *tactile TRFs* measured in each experimental condition. As previously reported, in the behavioral preliminary experiment, we measured participants’ ability to discriminate by sounds whether the experimenter was touching with the brush his own hand. However, we could not exclude the possibility that sounds could be informative below threshold. Similarly, while participants could not see their hands, nor the experimenters’, we ensured that no other visual cue could lead to spurious entrainment in synchrony with the tactile one. To this end, cluster-based permutations were performed between *tactile TRFs* at each condition and the control condition *control TRF* (Maris & Oostenveld, 2007, Monte Carlo method 1000 permutations, the cluster-based p-value was 0.025, accounting for two-tailed testing) across 0 to 400 ms time-lags and all electrodes.

We investigated whether the tactile TRF associated with the different fingers could allow us to measure a somatotopy of the brain response with the EEG. We evaluated the possibility of measuring topographical differences regarding digit lateralization (i.e., right pinkie vs. left pinkie and right thumb vs. left thumb) and digit representation within each hand (i.e., right thumb vs. right pinkie and left thumb vs. left pinkie). We normalized the TRFs associated with each condition computing the z-score between all electrodes of each experimental TRF value for each subject and time-point (each finger of each hand) to avoid that amplitude differences between them could drive spurious topographical effects (e.g., Urbach & Kutas, 2002). Once again, a series of cluster-based permutation tests were performed between tactile TRFs between experimental conditions (e.g., left vs. right thumb or right pinkie vs. right thumb; Maris & Oostenveld, 2007, Monte Carlo method 1000 permutations, the cluster-based p-value was 0.025, accounting for two-tailed testing) across 50 to 400 ms time-lags and all electrodes. The testing between conditions was performed starting from 50 ms since differences between the experimental conditions and their null counterparts only emerged from 50 ms of time lag onward (see Results: 3.2 EEG data analysis). Crucially, we performed our analysis on the whole window of significance. However, we expected topographical differences in the early phases of the TRFs (short timescales) since this early activity is usually associated with early somatosensory cortex activity (Houzè et al., 2011; Nierula et al., 2013), where finger somatotopy is more evident (Houzè et al., 2011; Nierula et al., 2013).

For each statistical contrast, we computed the unbiased estimate of Cohen’s d to quantify the magnitude of the effect. Regarding the contrast between the TRF and the *null* TRF or the *control* TRF, we employed the same cluster of electrodes in which the TRF was greater in amplitude (electrodes [E22, E25, E26, E27, E28] for right stimulation and electrodes [E42, E45, E46, E48, E49] for left stimulation). Conversely, we followed a data-driven approach for each experimental contrast between conditions. Cohen’s d was computed across all electrodes pertaining to each significant cluster and its time points. Confidence intervals were computed with a bootstrap method with 1000 permutations. We acknowledge that the effect could be inflated since they are computed on the same sample.

Lastly, we evaluated the consistency of the somatosensory entrainment, investigating the minimum tactile stimulation time necessary to obtain a reliable tactile TRF greater than the *null TRF*. The spatiotemporal profiles of each experimental condition (tactile TRF) were contrasted to a null effect (null TRF). Crucially, we built eight different TRF models starting from 2 and reaching 9 minutes of stimulation. After evaluating the normality of the distribution, a series of paired t-tests were performed between each model tactile TRF and the corresponding *null TRF*, using as critical alpha 0.006 (Bonferroni corrected, eight multiple tests).

## 3. Results

### 3.1 Behavioral data analysis

The behavioral data collected in the preliminary experiment resulted in a percentage of hit and omission that entered a paired t-test, crucially hit and omission did not differ (mean ± SE: Hit 0.51 ± 0.01; Omission 0.49 ± 0.01; p-value = 0.70). Subsequently, the d’, a measure of sensitivity, was computed to assess if participants were able to distinguish trials in which the stimulation was performed compared to when it was not performed, considering that a d’ near to 0 indicates absence of sensitivity (d’ = 1.6494e-15).

### 3.2 EEG data analysis

Two different sets of cluster-based permutated paired t-tests were performed. A preliminary analysis contrasting each tactile neural tracking versus the correspondent null effect to ensure a reliable statistical existence of the tactile TRF. A secondary analysis contrasting each experimental condition TRF versus the control condition, in which the experimenter was stimulating their own hand hidden from participants’ view. This further control analysis was performed to prove that tactile TRF could not be explained by spurious auditory or visual information.

#### Tactile entrainment vs. Null

As a first analysis, within each experimental condition (i.e., left thumb, left pinkie, right thumb, and right pinkie), the tactile TRF was contrasted with a corresponding *null TRF* at the whole brain level to capture the clusters of significant electrodes within the 0-400 ms time-lags. A clear tactile TRF emerged in each experimental condition, and this response was statistically different from the *null TRF*. For each experimental condition, one positive cluster (p_clust_ < 0.05) and one negative cluster (p_clust_ < 0.05) were identified on the contralateral central electrodes. Specifically, our results highlighted a central and contralateral positive response (50-170 ms; left pinkie: d=1.88, 95th confidence interval (CI95)=1.32 – 2.43; right pinkie: d=1.39, CI95=0.79 – 1.91; left thumb: d=1.4, CI95=0.88 – 2.02; right thumb: d=1.53, CI95=0.93 – 2.08) peaking around 140 ms, followed by central bilateral negativity (200-300 ms; left pinkie: d=-2.06, CI95=-2.85 – -1.36; right pinkie: d=-2.15, CI95=-2.69 – -1.6; left thumb: d=-1.34, CI95=-1.98 – -0.57; right thumb: d=-2.02, CI95=-2.47 – -1.53) peaking around 245 ms (see Figure 2).

**Fig. 2.**
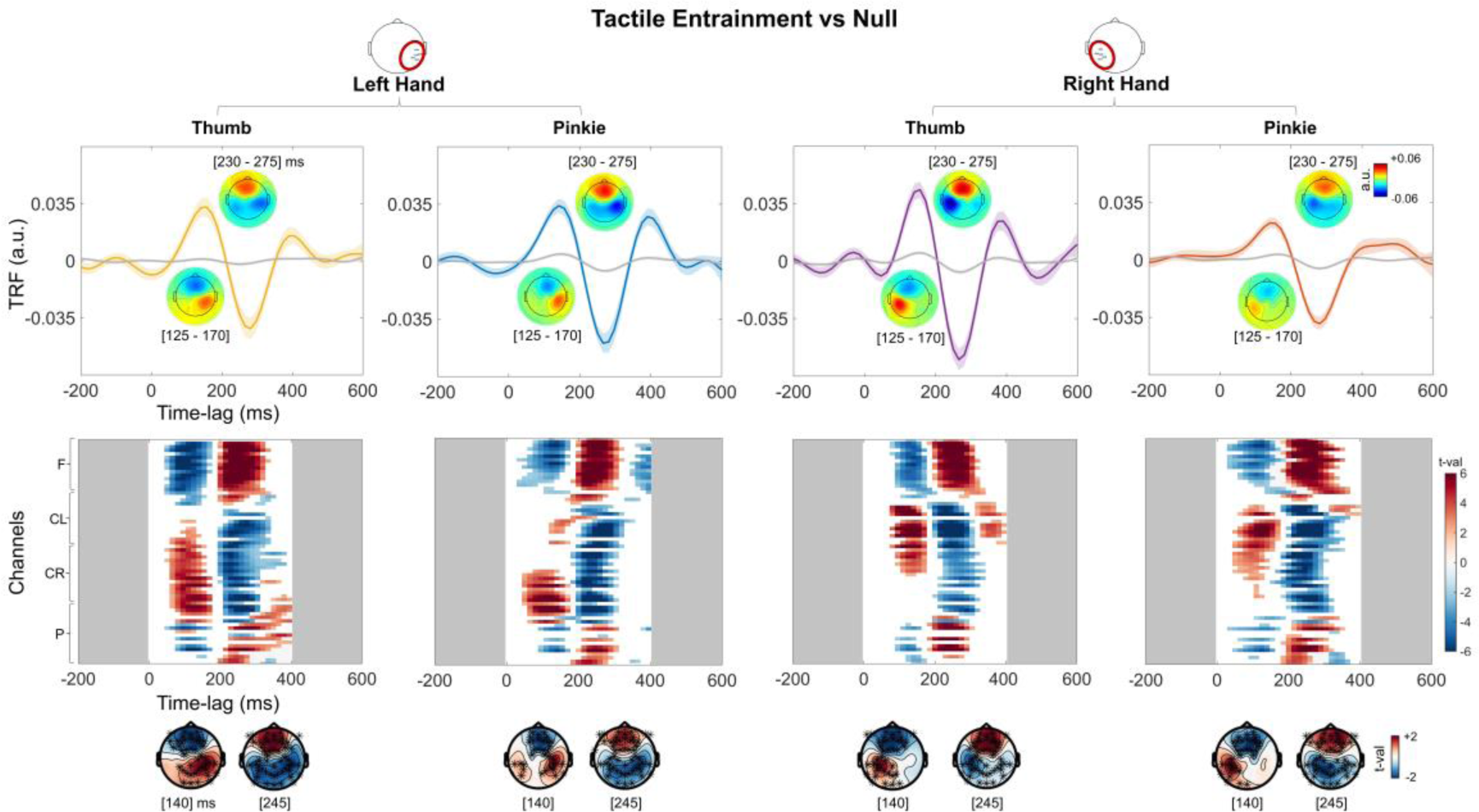
Tactile TRFs vs. *null TRFs*. Upper panel: the TRF corresponding to each experimental condition (colored lines) and the *null* TRF (in light grey). The *null TRF* is obtained by fitting the model on randomly mismatched pairs of stimulations (sound envelope of touch) and EEG responses. The TRFs and the null-TRFs represent here the average across central contralateral electrodes with respect to the finger stimulated. For all finger stimulations, the topographies reveal a first positive central contralateral activity (∼140ms) followed by a second negative bilateral response (∼245ms). Lower panel: results of the cluster-based permutation tests between the experimental TRF and null-TRF, across 0 to 400 ms time-lags and all sensors. For each experimental condition, a first positive response (p_clust_ < 0.05) and a second negative response (p_clust_ < 0.05) emerged.

#### Tactile entrainment vs. control

Subsequently, the *tactile TRF of each condition* was contrasted with the *control TRF* (of the control condition). A clear tactile response function was significant when controlling for auditory and visual spurious activity. Each experimental condition differed significantly from the control condition TRF, revealing one positive response at the central contralateral cluster (p_clust_ < 0.05) and a second negative response (p_clust_ < 0.05; See Figure 3). Critically, also with this contrast, two defined peaks emerged. Their statistical significance, topographical and temporal distribution were strictly similar to the previous one, with a central contralateral positive response (50-170 ms; left pinkie: d=1.31, CI95=0.57 – 1.78; right pinkie: d=0.68, CI95=0.24 – 1.12; left thumb: d=0.95, CI95=0.34 – 1.50; right thumb: d=0.97, CI95=0.39 – 1.46) peaking around 140 ms, followed by central bilateral negativity (200-300 ms; left pinkie: d=-1.66, CI95=-2.24 – -0.88; right pinkie: d=-1.14, CI95=-1.61 – - 0.58; left thumb: d=-1.04, CI95=-1.72 – -0.25; right thumb: d=-1.5, CI95=-1.98 – -0.86) peaking around 245 ms (see Figure 3).

**Fig. 3.**
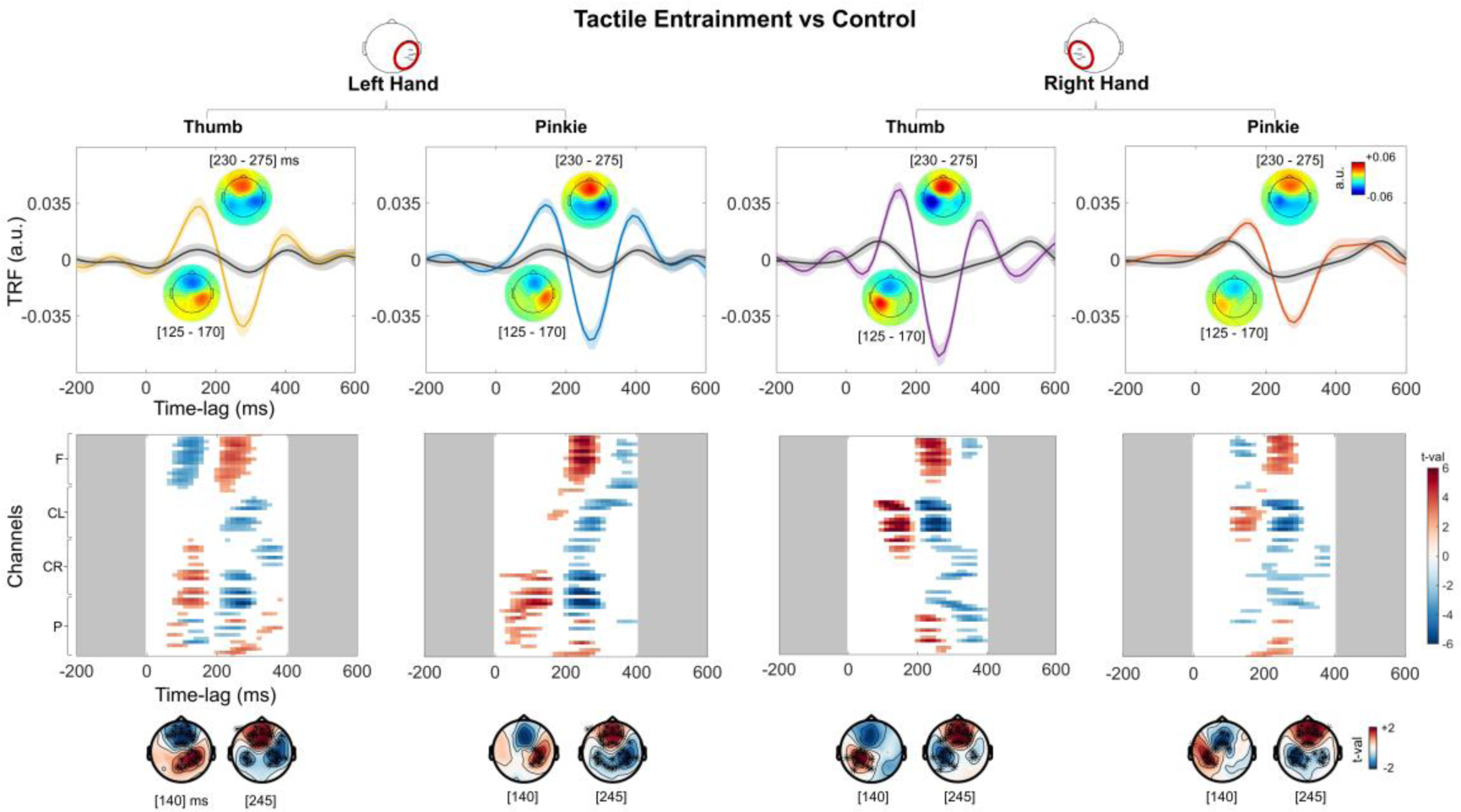
Tactile TRFs vs. *Control TRFs*. Upper panel: the TRF corresponding to each experimental condition (coloured lines, same TRFs reported in Fig. 2) and the *control* TRF (in black). The *control TRF* is obtained by keeping visual and auditory spurious activity constant but without the somatosensory component of the stimulation (i.e., the experimenter’s left thumb is stimulated and not the participant’s). The tactile TRFs and the control-TRFs represent here the average across central contralateral electrodes with respect to the finger stimulated. The topographies reveal a first positive contralateral peak (∼140ms) followed by a second negative bilateral response (∼245ms). Lower panel: results of the cluster-based permutation tests between the experimental TRF and *control* TRF, across 0 to 400 ms time-lags and all sensors. For each experimental condition, a positive response (p_clust_ < 0.05) and a second negative response (p_clust_ < 0.05) emerged.

#### Digit lateralization

To investigate digit lateralization, we compared the normalized topographies performing specific contrasts between the TRFs associated with the fingers across the hands. We contrasted the left thumb with the right thumb experimental TRFs (See Figure 4). This resulted in three statistically significant clusters. Specifically, our results highlighted an early phase central bilateral difference (50-185 ms, p_clust_ < 0.05, d=-1.55, CI95=-2.12 – -0.78) peaking at 110ms, a subsequent central bilateral difference (200-275 ms, p_clust_ < 0.05, d=0.10, CI95=-0.33 – 0.63) peaking at 230 ms, and a final bilateral difference (300-365 ms p_clust_ < 0.05, d=0.08, CI95=-0.30 – 0.46) peaking at 350 ms. Then, we contrasted the left pinkie vs the left thumb, resulting in two significant clusters. Specifically, our results highlighted a first central bilateral difference (50-170 ms, p_clust_ < 0.05, d=0.15, CI95=-0.39 – 0.65) peaking at 110 ms and a subsequent negative differential response (300-395 ms, p_clust_ < 0.05, d=-0.94, CI95=-1.46 – -0.46) peaking around 350 ms.

**Fig. 4.**
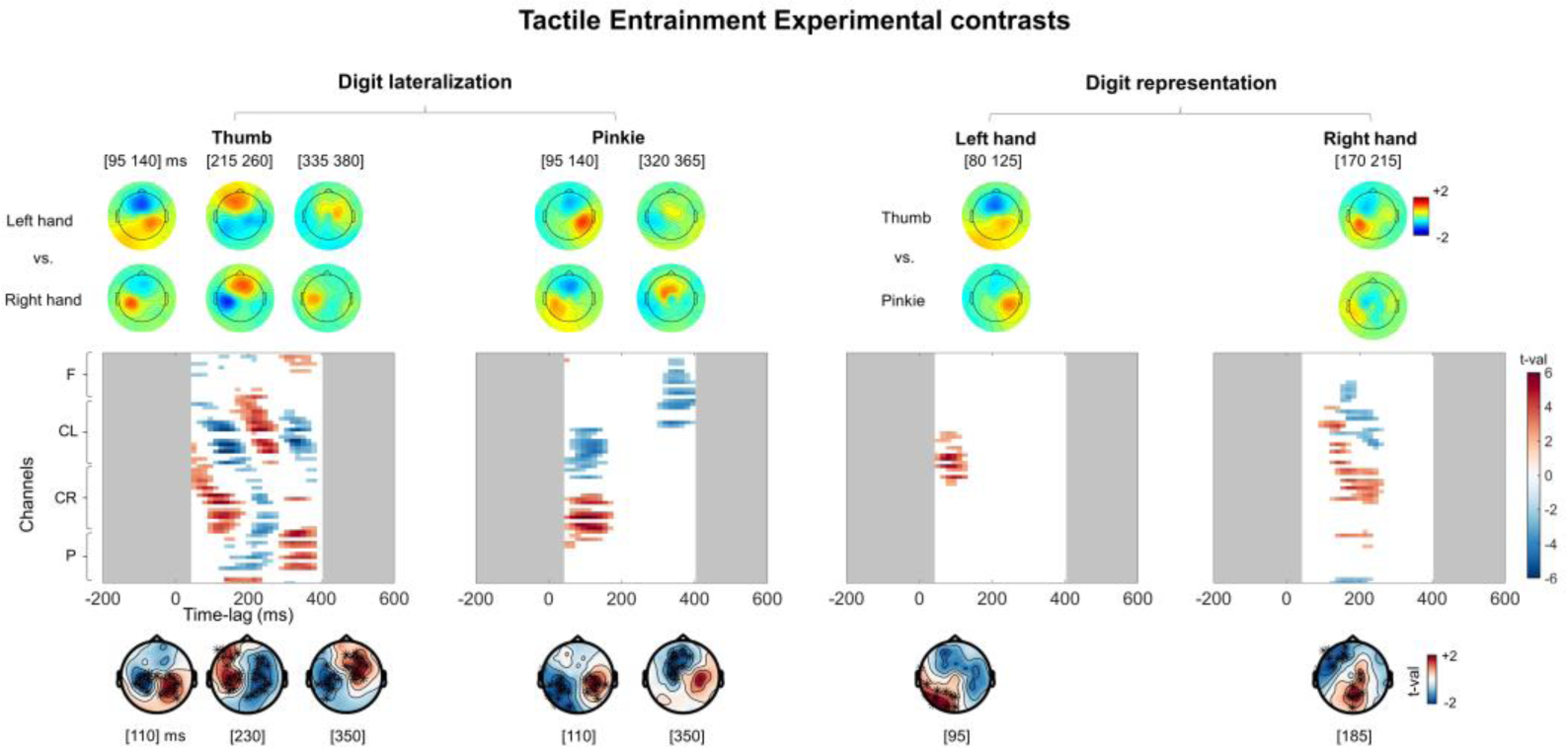
Contrasts between hands and between fingers. Results of the cluster-based permutation paired tests between the normalized TRFs for digit lateralization (left thumb vs. right thumb and left pinkie vs. right pinkie) and, for digit representation (left thumb vs. left pinkie and right thumb vs. right pinkie), across 50 to 400 ms time-lags and all sensors. Regarding the digit lateralization (left panel), for the left thumb vs. right thumb contrast, and for the left pinkie vs. right pinkie contrast, results highlighted topographical differences between the TRFs associated with fingers across hands (all p_clust_ < 0.05). Regarding digit representation, contrasting the left thumb vs. left pinkie and the right thumb vs. left thumb showed topographies differences across the TRFs associated with the fingers of each hand (all p_clust_ < 0.05). Results suggest that the tactile entrainment can be used to successfully measure the somatotopy of finger representation.

#### Digit representation

We investigated whether the topographies of the tactile TRF were sensitive to each finger within the left and right hands. We contrasted normalized topographies of the TRFs associated with the left thumb and the left pinkie. This resulted in a significant early phase cluster (50 – 125 ms p_clust_ < 0.05, d=1.10, CI95=0.69 – 1.54) peaking around 95 ms. Then, we contrasted the right thumb vs. right pinkie normalized topographies. This resulted in a protracted, statistically significant cluster (95 – 260 ms, p_clust_ < 0.05, d=0.26, CI95=-0.42 – 0.86) peaking around 185 ms.

#### Minimum data to measure tactile entrainment

Eventually, we contrasted tactile TRF and *null TRF* estimated by eight different models obtained using different lengths of tactile stimulation (2-9 minutes). Crucially, the contrast between tactile TRF and null-TRF was significant for each experimental condition, starting from 3 minutes of tactile stimulation (p-values< critical alpha: 0.0065).

## 4. Discussion

Neural tracking is an essential mechanism subserving an efficient interaction with the external world. It was reliably measured in the visual (Bourguignon et al., 2020; Hauswald et al., 2018; King et al., 2016; Spaak et al., 2014) and auditory (Di Liberto et al., 2015; Keitel et al., 2017; Kösem et al., 2018; Zoefel & VanRullen, 2016; Zuk et al., 2021) systems following continuous stimulation. Here, we extended the measurement of neural tracking to the somatosensory domain, employing a continuous and naturalistic tactile stimulation of the hands. Notably, each participant received unique and random stimulation with a series of gentle brushes. We modeled the sound generated by tactile stimulation, namely the recording of the friction between the skin and the miniaturized microphone inserted in a brush. In defining the tactile envelope, the frequency range with the greater energy (i.e., 0-6 Hz, see Figure 1) was chosen to have a comprehensive descriptor of the data with maximal information.

First, we estimated tactile neural tracking at the individual level employing a temporal response function (TRF). Critically, this is done with a leave-one-out cross-validation, validating each model at the single-subject level, considering the intrinsic variability of each participant and each one of the stimulations performed. Second, we estimated a group-level TRF and tested its existence by comparing it with a *null* TRF obtained by the random permutation of the data. Third, we calculated the minimum amount of data to estimate a robust group level TRF associated with continuous somatosensory stimulation for each finger; three minutes of passive tactile stimulation were sufficient to obtain a group level TRF significantly different from the *null TRF*, calculated with the same amount of data. Fourthly, contrasting the TRF obtained by the stimulation of single fingers of the hand vs. a *control TRF*, in which we controlled for possible auditory and visual confounds, a clear tactile response function emerged, confirming the measurement of a pure tactile entrainment with continuous tactile stimulation. Finally, regarding digit lateralization, contrasting the TRFs associated with left and right fingers, we found topographical changes depending on hand lateralization for both fingers at several time lags. Similarly, the spatiotemporal profiles of the TRFs allowed the dissociation of digit representation depending on the specific finger stimulation (i.e., thumb vs. pinkie) both for the left and right hand.

Only few preliminary observations employed continuous tactile stimulation while recording the EEG (Chota et al., 2023; Fu & Riecke, 2023). Most of the studies investigating the somatosensory system still rely on somatosensory evoked potentials (SEPs) and averaged responses across several trials (Fossataro et al., 2023a; Hogendoorn et al., 2015; Macerollo et al., 2018; Pyasik et al., 2021; Ronga et al., 2021). Among the few studies which investigated neural responses to continuous tactile input, Chota and colleagues investigated the role of phase-locked β bursts following a vibrotactile artificial stimulation determined by the somatosensory system resonance properties. Crucially, also when the somatosensory input was mapped continuously (Chota et al., 2023; Fu & Riecke, 2023), these studies measured steady-state somatosensory evoked potentials (SSSEPs) that required a highly standardized series of events with a specific predefined frequency of stimulation. Here, we measured how the brain encodes dynamic somatosensory events, as in tactile interactions. Despite employing unique and different stimulations for each condition and participant, we obtained a robust and reliable tactile TRF. This result was possible because for everyone, the tactile TRF was estimated by modeling the envelope of the tactile information describing the time series and the dynamics of tactile events. Critically, each stimulation differed randomly in several parameters, like the onset/offset of each single brush, the duration and intensity of each brush, and the locus of stimulation on the finger. In other words, the variability of the stimulation was modeled and used to gain generalizability of findings. We focused our analysis on slow tactile somatosensory stimulations that the sound envelope could convey below 6 Hz. However, continuous tactile stimuli are potentially richer, and other features could be analyzed in the future, such as direction of motion, speed, and texture.

To assess the reliability of this protocol, it was important to evaluate the consistency in terms of timing and topography with previous studies employing somatosensory stimuli. Noteworthy, by measuring the encoding of finger brushing we modelled slow tactile events (sound envelope below 6 Hz) on slow rhythmic brain activity (below 6 Hz). This prevented us from measuring faster tactile response functions. Classical clinical SEPs studies highlight early components through the stimulation of the median nerve, leading to a complex chain of early responses (i.e., P14, N20, P22, N24 and N30) starting before 30 ms after the stimulus onset (Valeriani et al., 2001; Vibell et al., 2023). However, these early subcortical components are visible following the average across several hundred trials (Vibell et al., 2023) and are not measurable following skin stimulations (Zeller et al., 2015). Conversely, the cortical SEPs generally include the N25, P45, N80, P1/P100, N1/N140, P2/P200, N2 (about 250 ms after touch onset), and P3 components, with early contralateral topography followed by later bilateral responses (Cardini et al., 2012; Nuwer, 1998; Sambo & Forster, 2008). Our results are in line with well-established cortical SEPs and highlighted in the TRF a clear first contralateral positive response at central electrodes which started at about 50 ms of lag; this response was compatible with early somatosensory processing components highlighted in ERP studies (Shimojo et al., 2000; Zeller et al., 2015). This response peaked at around 160 ms and was followed by a central bilateral negativity peaking at about 260 ms. Both the timing and topography of these responses align with the SEPs literature (Eimer & Forster, 2003; Sambo & Forster, 2008; Shimojo et al., 2000; Zeller et al., 2015; Vibell et al., 2023), suggesting that the method unveiled consistent neural tracking of tactile stimulation.

It is important to stress that the encoding model of the TRF was statistically tested with a data driven approach, controlling for the multiple comparisons problem and contrasting TRFs with null effects (Maris & Oostenveld, 2007). In canonical ERPs research, the brain responses are usually averaged and contrasted between experimental conditions. Here, the approach provides control by variability since the entire tactile information is modeled and the response of interest can be objectively identified in time (time lags) and space (scalp) domains, strictly when it exceeds the estimation of noise or control conditions, such as the *null TRF* and the *control TRF*. In addition to its effectiveness, this paradigm strength also relies on the limited amount of time required to obtain a reliable tactile response function. Indeed, despite idiosyncratic stimulations (every trial was different for every finger and participant), we measured a significant and stable TRF already with three minutes of stimulation. This indication can drastically shorten the experimental paradigms employing tactile stimulation, allowing the contrast of several experimental conditions within a relatively short experimental session, especially in populations such as infants (Ronga et al., 2021), people with autistic spectrum disorders (Voos et al., 2013) or clinical and psychiatric population (Keizer et al., 2022).

Results clearly demonstrated a pure tactile TRF. In addition to the contrast with the null effect, the tactile TRF was contrasted with the *control TRF*. The control condition was identical to the experimental one regarding auditory and visual cues, such as the noise associated with brushing or the experimenter’s movements. This allowed to control for potential auditory and visual inputs collinear to the tactile stimulation. However, since the experimenter stimulated his own hands, the participants did not have any tactile stimulation. Therefore, contrasting the tactile TRF with the *control TRF* ensured estimating actual tactile entrainment, avoiding spurious effects associated with other input processing.

Lastly, our methodology successfully differentiated the topography of a somatosensory response to a continuous and naturalistic stimulation on the thumb or the pinkie, independently of the stimulated hand. Whereas a digitotopy of the hand has been described employing fMRI (O’Neill et al., 2020; Sanchez-Panchuelo et al., 2012; Sanchez-Panchuelo et al., 2010) or MEG (Sun et al., 2021; Nakamura et al., 1998), only few EEG studies dissociated the neural response at the level of the stimulated finger. Houzè and colleagues (2011) showed a difference between SEP sources of the N20/P20 component of the thumb and pinkie. A subsequent study (Nierula et al., 2013) confirmed Houzè and colleagues’ results in the SEPs domain (Nierula et al., 2013). Crucially, both studies employed SEP elicited by electrical stimulation, clearly a non-ecological type of tactile stimulation.

This study opens new venues for future work employing similar analytical approaches to evaluate different populations as well as experimental manipulations of somatosensory processing. With an ontogenetic perspective, it will be essential to investigate how the tactile entrainment emerges by describing its developmental trajectory, opening the possibility of using this measure as a marker of typical and atypical development. Studies in the animal domain could provide the phylogenesis of this tactile response function. This method could additionally help characterize and further distinguish the discriminative touch from the affective touch (McGlone et al., 2014; Singh et al., 2014) or tickle, a specific somatosensory experience that has intrinsic value as a mediator of the social aspect of touch both in the phylogenetic and ontogenetic development, employing and evaluating online and real-life tactile interactions among individuals. Due to its intrinsic properties, it is not easy to assess how the human brain elaborates affective touch (i.e., a continuous tactile stimulation with a preferential velocity of around 1-10 cm/s that activates the CT-fibers). Indeed, it is impossible to reproduce this affective component of touch with highly standardized laboratory stimuli, while it is possible with a naturalistic and continuous stimulation. Currently, most EEG studies investigate the early and late components of late positive potential (LPP) or modulations of brain activity in specific frequency bands (McGlone et al., 2014; Portnova et al., 2020; Schirmer & Gunter, 2017; Singh et al., 2014; von Mohr et al., 2018). The tactile TRF should be able to highlight and characterize when and how affective touch is processed in the brain and its difference from discriminative touch simply by changing stimulation properties. Finally, the current approach could be used to stimulate different body parts and evaluate eventual differences in neural tracking associated with different body districts. Our results pave the way for a more efficient and naturalistic study of the somatosensory system, evaluating how it works in situations more affine at how the brain elaborates tactile information in real-time daily interactions.

## References

Bourguignon, M., Baart, M., Kapnoula, E. C., & Molinaro, N. (2020). Lip-reading enables the brain to synthesize auditory features of unknown silent speech. Journal of Neuroscience, 40(5), 1053–1065. 10.1523/JNEUROSCI.1101-19.2019

Campos Viola, F., Thorne, J., Edmonds, B., Schneider, T., Eichele, T., & Debener, S. (2009). Semi-automatic identification of independent components representing EEG artifact. Clinical Neurophysiology, 120(5), 868–877. 10.1016/j.clinph.2009.01.015

Cardini, F., Longo, M. R., Driver, J., & Haggard, P. (2012). Rapid enhancement of touch from non-informative vision of the hand. Neuropsychologia, 50(8), 1954– 1960. 10.1016/j.neuropsychologia.2012.04.020

Chota, S., VanRullen, R., & Gulbinaite, R. (2023). Random Tactile Noise Stimulation Reveals Beta-Rhythmic Impulse Response Function of the Somatosensory System. Journal of Neuroscience, 43(17), 3107–3119. 10.1523/JNEUROSCI.1758-22.2023

Crosse, M. J., Di Liberto, G. M., Bednar, A., & Lalor, E. C. (2016). The multivariate temporal response function (mTRF) toolbox: A MATLAB toolbox for relating neural signals to continuous stimuli. Frontiers in Human Neuroscience, 10(NOV2016). 10.3389/fnhum.2016.00604

Crosse, M. J., Zuk, N. J., Di Liberto, G. M., Nidiffer, A. R., Molholm, S., & Lalor, E. C. (2021). Linear Modeling of Neurophysiological Responses to Speech and Other Continuous Stimuli: Methodological Considerations for Applied Research. In Frontiers in Neuroscience (Vol. 15). Frontiers Media S.A. 10.3389/fnins.2021.705621

Delorme, A., & Makeig, S. (2004). EEGLAB: An open source toolbox for analysis of single-trial EEG dynamics including independent component analysis. Journal of Neuroscience Methods, 134(1), 9–21. 10.1016/j.jneumeth.2003.10.009

Di Liberto, G. M., O’Sullivan, J. A., & Lalor, E. C. (2015). Low-frequency cortical entrainment to speech reflects phoneme-level processing. Current Biology, 25(19), 2457–2465. 10.1016/j.cub.2015.08.030

Eimer, M., & Forster, B. (2003). The spatial distribution of attentional selectivity in touch: Evidence from somatosensory ERP components. Clinical Neurophysiology, 114(7), 1298–1306. 10.1016/S1388-2457(03)00107-X

Fossataro, C., Galigani, M., Rossi Sebastiano, A., Bruno, V., Ronga, I., & Garbarini, F. (2023). Spatial proximity to others induces plastic changes in the neural representation of the peripersonal space. IScience, 26(1). 10.1016/j.isci.2022.105879

Fu, X., & Riecke, L. (2023). Effects of continuous tactile stimulation on auditory-evoked cortical responses depend on the audio-tactile phase. NeuroImage, 274. 10.1016/j.neuroimage.2023.120140

Galigani, M., Ronga, I., Fossataro, C., Bruno, V., Castellani, N., Rossi Sebastiano, A., Forster, B., & Garbarini, F. (2021). Like the back of my hand: Visual ERPs reveal a specific change detection mechanism for the bodily self. Cortex, 134, 239–252. 10.1016/j.cortex.2020.10.014

Haufe, S., Meinecke, F., Görgen, K., Dähne, S., Haynes, J. D., Blankertz, B., & Bießmann, F. (2014). On the interpretation of weight vectors of linear models in multivariate neuroimaging. NeuroImage, 87, 96–110. 10.1016/j.neuroimage.2013.10.067

Hauswald, A., Lithari, C., Collignon, O., Leonardelli, E., & Weisz, N. (2018). A Visual Cortical Network for Deriving Phonological Information from Intelligible Lip Movements. Current Biology, 28(9), 1453–1459.e3. 10.1016/j.cub.2018.03.044

Hogendoorn, H., Kammers, M., Haggard, P., & Verstraten, F. (2015). Self-touch modulates the somatosensory evoked P100. Experimental Brain Research, 233(10), 2845–2858. 10.1007/s00221-015-4355-0

Houzé, B., Perchet, C., Magnin, M., & Garcia-Larrea, L. (2011). Cortical representation of the human hand assessed by two levels of high-resolution EEG recordings. Human Brain Mapping, 32(11), 1894–1904. 10.1002/hbm.21155

Jessen, S., Fiedler, L., Münte, T. F., & Obleser, J. (2019). Quantifying the individual auditory and visual brain response in 7-month-old infants watching a brief cartoon movie. NeuroImage, 202. 10.1016/j.neuroimage.2019.116060

Jung, T., Makeig, S., Humphries, C., Lee, T., McKeown, M. J., Iragui, V., & Sejnowski, T. J. (2000). Removing electroencephalographic artifacts by blind source separation. Psychophysiology, 37(2), 163–178. 10.1111/1469-8986.3720163

Keitel, A., Ince, R. A. A., Gross, J., & Kayser, C. (2017). Auditory cortical delta-entrainment interacts with oscillatory power in multiple fronto-parietal networks. NeuroImage, 147, 32–42. 10.1016/j.neuroimage.2016.11.062

Keizer, A., Heijman, J. O., & Dijkerman, H. C. (2022). Do transdiagnostic factors influence affective touch perception in psychiatric populations? In Current Opinion in Behavioral Sciences (Vol. 43, pp. 125–130). Elsevier Ltd. 10.1016/j.cobeha.2021.09.006

King, A. J., Park, H., Kayser, C., Thut, G., & Gross, J. (2016). Lip movements entrain the observers’ low-frequency brain oscillations to facilitate speech intelligibility. 10.7554/eLife.14521.001

Kösem, A., Bosker, H. R., Takashima, A., Meyer, A., Jensen, O., & Hagoort, P. (2018). Neural Entrainment Determines the Words We Hear. Current Biology, 28(18), 2867–2875.e3. 10.1016/j.cub.2018.07.023

Lakatos, P., Gross, J., & Thut, G. (2019). A New Unifying Account of the Roles of Neuronal Entrainment. In Current Biology (Vol. 29, Issue 18, pp. R890–R905). Cell Press. 10.1016/j.cub.2019.07.075

Maallo, A. M. S., Duvernoy, B., Olausson, H., & McIntyre, S. (2022). Naturalistic stimuli in touch research. In Current Opinion in Neurobiology (Vol. 75). Elsevier Ltd. 10.1016/j.conb.2022.102570

Macerollo, A., Brown, M. J. N., Kilner, J. M., & Chen, R. (2018). Neurophysiological Changes Measured Using Somatosensory Evoked Potentials. In Trends in Neurosciences (Vol. 41, Issue 5, pp. 294–310). Elsevier Ltd. 10.1016/j.tins.2018.02.007

Maris, E., & Oostenveld, R. (2007). Nonparametric statistical testing of EEG- and MEG-data. Journal of Neuroscience Methods, 164(1), 177–190. 10.1016/j.jneumeth.2007.03.024

McGlone, F., Wessberg, J., & Olausson, H. (2014). Discriminative and Affective Touch: Sensing and Feeling. In Neuron (Vol. 82, Issue 4, pp. 737–755). Cell Press. 10.1016/j.neuron.2014.05.001

Mirkovic, B., Debener, S., Jaeger, M., & De Vos, M. (2015). Decoding the attended speech stream with multi-channel EEG: Implications for online, daily-life applications. Journal of Neural Engineering, 12(4). 10.1088/1741-2560/12/4/046007

Morales, S., & Bowers, M. E. (2022). Time-frequency analysis methods and their application in developmental EEG data. Developmental Cognitive Neuroscience, 54. 10.1016/j.dcn.2022.101067

Nakamura, A., Yamada, T., Goto, A., Kato, T., Ito, K., Abe, Y., Kachi, T., & Kakigi, R. (1998). Somatosensory Homunculus as Drawn by MEG.

Nierula, B., Hohlefeld, F. U., Curio, G., & Nikulin, V. V. (2013). No somatotopy of sensorimotor alpha-oscillation responses to differential finger stimulation. NeuroImage, 76, 294–303. 10.1016/j.neuroimage.2013.03.025

Nuwer, M. R. (1998). Fundamentals of evoked potentials and common clinical applications today.

Obleser, J., & Kayser, C. (2019). Neural Entrainment and Attentional Selection in the Listening Brain. In Trends in Cognitive Sciences (Vol. 23, Issue 11, pp. 913–926). Elsevier Ltd. 10.1016/j.tics.2019.08.004

O’Neill, G. C., Sengupta, A., Asghar, M., Barratt, E. L., Besle, J., Schluppeck, D., Francis, S. T., & Sanchez Panchuelo, R. M. (2020). A probabilistic atlas of finger dominance in the primary somatosensory cortex. NeuroImage, 217. 10.1016/j.neuroimage.2020.116880

Pion-Tonachini, L., Kreutz-Delgado, K., & Makeig, S. (2019). ICLabel: An automated electroencephalographic independent component classifier, dataset, and website. NeuroImage, 198, 181–197. 10.1016/j.neuroimage.2019.05.026

Plass, J., Brang, D., Suzuki, S., & Grabowecky, M. (2020). Vision perceptually restores auditory spectral dynamics in speech. 117(29), 16920–16927. 10.1073/pnas.2002887117/-/DCSupplemental

Portnova, G. V., Proskurnina, E. V., Sokolova, S. V., Skorokhodov, I. V., & Varlamov, A. A. (2020). Perceived pleasantness of gentle touch in healthy individuals is related to salivary oxytocin response and EEG markers of arousal. Experimental Brain Research, 238(10), 2257–2268. 10.1007/s00221-020-05891-y

Pyasik, M., Ronga, I., Burin, D., Salatino, A., Sarasso, P., Garbarini, F., Ricci, R., & Pia, L. (2021). I’m a believer: Illusory self-generated touch elicits sensory attenuation and somatosensory evoked potentials similar to the real self-touch. NeuroImage, 229. 10.1016/j.neuroimage.2021.117727

Ronga, I., Galigani, M., Bruno, V., Castellani, N., Rossi Sebastiano, A., Valentini, E., Fossataro, C., Neppi-Modona, M., & Garbarini, F. (2021). Seeming confines: Electrophysiological evidence of peripersonal space remapping following tool-use in humans. Cortex, 144, 133–150. 10.1016/j.cortex.2021.08.004

Ronga, I., Galigani, M., Bruno, V., Noel, J.-P., Gazzin, A., Perathoner, C., Serino, A., & Garbarini, F. (2021). Spatial tuning of electrophysiological responses to multisensory stimuli reveals a primitive coding of the body boundaries in newborns. PNAS. 10.1073/pnas.2024548118/-/DCSupplemental

Sambo, C. F., & Forster, B. (2008). An ERP Investigation on Visuotactile Interactions in Peripersonal and Extrapersonal Space: Evidence for the Spatial Rule.

Sanchez-Panchuelo, R. M., Besle, J., Beckett, A., Bowtell, R., Schluppeck, D., & Francis, S. (2012). Within-digit functional parcellation of brodmann areas of the human primary somatosensory cortex using functional magnetic resonance imaging at 7 tesla. Journal of Neuroscience, 32(45), 15815–15822. 10.1523/JNEUROSCI.2501-12.2012

Sanchez-Panchuelo, R. M., Francis, S., Bowtell, R., & Schluppeck, D. (2010). Mapping human somatosensory cortex in individual subjects with 7T functional MRI. Journal of Neurophysiology, 103(5), 2544–2556. 10.1152/jn.01017.2009

Schirmer, A., & Gunter, T. C. (2017). The right touch: Stroking of CT-innervated skin promotes vocal emotion processing. *Cognitive*, Affective and Behavioral Neuroscience, 17(6), 1129–1140. 10.3758/s13415-017-0537-5

Serino, A., & Haggard, P. (2010). Touch and the body. In Neuroscience and Biobehavioral Reviews (Vol. 34, Issue 2, pp. 224–236). 10.1016/j.neubiorev.2009.04.004

Shimojo, M., Svensson, P., Arendt-Nielsen, L., & Chen, A. C. N. (2000). Dynamic brain topography of somatosensory evoked potentials and equivalent dipoles in response to graded painful skin and muscle stimulation. Brain Topography, 13(1), 43–58. 10.1023/A:1007834319135

Singh, H., Bauer, M., Chowanski, W., Sui, Y., Atkinson, D., Baurley, S., Fry, M., Evans, J., & Bianchi-Berthouze, N. (2014). The brain’s response to pleasant touch: An EEG investigation of tactile caressing. Frontiers in Human Neuroscience, 8(November). 10.3389/fnhum.2014.00893

Spaak, E., de Lange, F. P., & Jensen, O. (2014). Local entrainment of alpha oscillations by visual stimuli causes cyclic modulation of perception. Journal of Neuroscience, 34(10), 3536–3544. 10.1523/JNEUROSCI.4385-13.2014

Sun, F., Zhang, G., Ren, L., Yu, T., Ren, Z., Gao, R., & Zhang, X. (2021). Functional organization of the human primary somatosensory cortex: A stereo-electroencephalography study. Clinical Neurophysiology, 132(2), 487–497. 10.1016/j.clinph.2020.11.032

Thézé, R., Giraud, A.-L., & Mégevand, P. (2020). The phase of cortical oscillations determines the perceptual fate of visual cues in naturalistic audiovisual speech. In Sci. Adv (Vol. 6). http://advances.sciencemag.org/

Valeriani, M., Le Pera, D., & Tonali, P. (2001). Characterizing somatosensory evoked potential sources with dipole models: Advantages and limitations. In Muscle and Nerve (Vol. 24, Issue 3, pp. 325–339). 10.1002/1097-4598(200103)24:3<25::AID-MUS1002>3.0.CO;2-0

von Mohr, M., Crowley, M. J., Walthall, J., Mayes, L. C., Pelphrey, K. A., & Rutherford, H. J. V. (2018). EEG captures affective touch: CT-optimal touch and neural oscillations. *Cognitive*, Affective and Behavioral Neuroscience, 18(1), 155–166. 10.3758/s13415-017-0560-6

Voos, A. C., Pelphrey, K. A., & Kaiser, M. D. (2013). Autistic traits are associated with diminished neural response to affective touch. Social Cognitive and Affective Neuroscience, 8(4), 378–386. 10.1093/scan/nss009

Wang, C., & Zhang, Q. (2021). Word frequency effect in written production: Evidence from ERPs and neural oscillations. Psychophysiology, 58(5). 10.1111/psyp.13775

Zeller, D., Litvak, V., Friston, K. J., & Classen, J. (2015). Sensory processing and the rubber hand Illusion—An evoked potentials study. Journal of Cognitive Neuroscience, 27(3), 573–582. 10.1162/jocn_a_00705

Zoefel, B., & VanRullen, R. (2016). EEG oscillations entrain their phase to high-level features of speech sound. NeuroImage, 124, 16–23. 10.1016/j.neuroimage.2015.08.054

Zuk, N. J., Murphy, J. W., Reilly, R. B., & Lalor, E. C. (2021). Envelope reconstruction of speech and music highlights stronger tracking of speech at low frequencies. PLoS Computational Biology, 17(9). 10.1371/journal.pcbi.1009358

